## Supplementary Tables for "The non-specific phospholipase C of common bean *PvNPC4* modulates roots and nodule development"

S1 Table. Primer sequences used for qPCR and plasmid construction.

| **Primers** | **Sequence 5´-3´** | **Usage** |
| --- | --- | --- |
| qFw-NPC4 | TCCTCAGTTCTTGACAAAGCG | *PvNPC4* transcript quantification in wild-type tissues |
| qRv-NPC4 | CATCACCACCAACCTCTCTTAG |  |
| qFw2-NPC4 | GCATTGTGAAGAAGGGAGATTGG | *PvNPC4* transcript quantification in transgenic roots |
| qRv2-NPC4 | GAGGACTTGTTCTTAGTGCCTC |  |
| qFw-PI-PLC4 | GGTGTTTAACAGTGGGTAACGGG | *PvPI-PLC4* transcript quantification in wild-type tissues |
| qRv-PI-PLC4 | GTGGATTCCGAGTTTACTTCAACC |  |
| EF1α-Fw | GGTCATTGGTCATGTCGACTCTGG | Transcript quantification |
| EF1α-Rv | GCACCCAGGCATACTTGAATGACC |  |
| β-tubulin-Fw | GGTGAGTGAGCAGTTCACTG | Transcript quantification |
| β-tubulin-Rv | CATCCTCATCTGCAGTTGC |  |
| PvEnod40qFw | AGT TTT GTT GGC AAG CAT CC | Transcript quantification |
| PvEnod40qRv | TAA GCA CAA GCA AAC TGT TG |  |
| PvNIN-qFw | GGGGATTCAGAGATTTGCAG | Transcript quantification |
| PvNIN-qRv | AACCCACTCTTGAGCATCGT |  |
| PvTML1a-qFw | ACA CTG GTT CTC CTG ATC CTG |  |
| PvTML1a-qRv | TTC AGG TCC ACA TCA CAG CAC |  |
| PvASL18a-qFw | AAG CAC AGC TGA TGC AGG TG |  |
| PvASL18a-qRv | TGG ATG GTT CAT TGG TTG TCC |  |
| PvASL18b-qFw | GCA GTA TGA ATG ATG GAA TGA GC |  |
| PvASL18b-qRv | TCA CAC TTC CTC ATC ATC CTG AG |  |
| FwRNAi-NPC4 | caccTCCTCAGTTCTTGACAAAGCG | RNAi fragment amplification |
| RvRNAi-NPC4 | CATCACCACCAACCTCTCTTAG |  |
| M13 rev | CAGGAAACAGCTATGAC | Verification of correct insertion in donor vector |
| qFw-NPC4 | TCCTCAGTTCTTGACAAAGCG |  |
| 35ptdT 9067 | GGAAGTTCATTTCATTTGGAGAGAAC | Verification of correct insertion in *PvNPC4*:RNAi vector |
| NWRKY-Rv | CTTCCGATCACGATTCATTGTC |  |
| NWRKY-Fw | GAAGTCTGAACAATTCTTGGGATTG |  |
| qFw-NPC4 | TCCTCAGTTCTTGACAAAGCG |  |

S2 Table. List of PLCs of *L. japonicus* and *M. truncatula*

| **Identifier** | **Protein name** | **Subfamily** |
| --- | --- | --- |
| Lj2g3v2256390.1 | LjPI-PLC1 | PI-PLC |
| Lj2g3v2256400.1 | LjPI-PLC2 |  |
| Lj2g3v2748690.1 | LjPI-PLC3 |  |
| Lj6g3v0521600.2 | LjPI-PLC4 |  |
| Lj6g3v0521670.1 | LjPI-PLC5 |  |
| Lj0g3v0130579.1 | LjPI-PLC6 |  |
| Lj1g3v1500500.1 | LjNPC1 | NPC |
| Lj6g3v0964670.1 | LjNPC2 |  |
| Lj0g3v0152469.1 | LjNPC3 |  |
| Lj0g3v0158969.1 | LjNPC4 |  |
| Lj0g3v0198969.1 | LjNPC5 |  |
| Lj0g3v0241399.1 | LjNPC6 |  |
| Medtr3g069280.1 | MtPI-PLC1 | PI-PLC |
| Medtr3g070560.1 | MtPI-PLC2 |  |
| Medtr3g070710.1 | MtPI-PLC3 |  |
| Medtr3g070720.1 | MtPI-PLC4 |  |
| Medtr5g071010.1 | MtPI-PLC5 |  |
| Medtr5g071040.1 | MtPI-PLC6 |  |
| Medtr5g082580.1 | MtPI-PLC7 |  |
| Medtr5g082620.1 | MtPI-PLC8 |  |
| Medtr5g082590.1 | MtPI-PLC9 |  |
| Medtr3g040680.1 | MtNPC1 | NPC |
| Medtr3g099120.2 | MtNPC2 |  |
| Medtr3g463760.1 | MtNPC3 |  |
| Medtr8g010590.1 | MtNPC4 |  |
| Medtr8g010690.1 | MtNPC5 |  |
| Medtr8g020860.1 | MtNPC6 |  |
| Glyma.02G257000.1 | GmPI-PLC1 | PI-PLC |
| Glyma.02G257100.1 | GmPI-PLC2 |  |
| Glyma.02G257200.1 | GmPI-PLC3 |  |
| Glyma.11G229900.1 | GmPI-PLC4 |  |
| Glyma.11G230000.1 | GmPI-PLC5 |  |
| Glyma.11G230100.1 | GmPI-PLC6 |  |
| Glyma.14G059200.1 | GmPI-PLC7 |  |
| Glyma.14G059300.1 | GmPI-PLC8 |  |
| Glyma.14G059400.1 | GmPI-PLC9 |  |
| Glyma.14G193800.1 | GmPI-PLC10 |  |
| Glyma.18G027100.1 | GmPI-PLC11 |  |
| Glyma.18G027200.1 | GmPI-PLC12 |  |
| Glyma.18G027300.1 | GmPI-PLC13 |  |
| Glyma.03G092800.1 | GmNPC1 | NPC |
| Glyma.04G196700.1 | GmNPC2 |  |
| Glyma.06G169100.1 | GmNPC3 |  |
| Glyma.11G178400.1 | GmNPC4 |  |
| Glyma.15G188700.1 | GmNPC5 |  |
| Glyma.16G081200.1 | GmNPC6 |  |
| Glyma.18G064000.1 | GmNPC7 |  |
| Glyma.18G064200.1 | GmNPC8 |  |
| Glyma.20G040600.1 | GmNPC9 |  |

S3 Table. List and classification of the 12 PvPLC

| **Gen identifier** | **Chromosome location** | **Subfamily** | **Gene name** |
| --- | --- | --- | --- |
| Phvul.001G243100.1 | 1 | PI-PLC | *PI-PvPLC1* |
| Phvul.001G243200.1 | 1 |  | *PI-PvPLC2* |
| Phvul.001G243300.1 | 1 |  | *PI-PvPLC3* |
| Phvul.008G224000.1 | 8 |  | *PI-PvPLC4* |
| Phvul.008G224100.2 | 8 |  | *PI-PvPLC5* |
| Phvul.008G224200.1 | 8 |  | *PI-PvPLC6* |
| Phvul.008G261100.1 | 8 |  | *PI-PvPLC7* |
| Phvul.006G014100.1 | 6 | NPC | *PvNPC1* |
| Phvul.006G066600.1 | 6 |  | *PvNPC2* |
| Phvul.009G165500.1 | 9 |  | *PvNPC3* |
| Phvul.010G033400.1 | 10 |  | *PvNPC4* |
| Phvul.L001661.1 | Scaffold_293 |  | *PvNPC5* |

S4 Table. Physicochemical properties of PvPLCs

| **Name** | **Length**  **(aa)** | **MW**  **(kDa)** | **IP** | **I. index** | **GRAVI** | **Subcellular localization** |
| --- | --- | --- | --- | --- | --- | --- |
| PI-PvPLC1 | 636 | 72.54 | 9.23 | 42.15 | -0.615 | Nuclear |
| PI-PvPLC2 | 550 | 62.41 | 6.3 | 42.54 | -0.471 | Cytoplasmic |
| PI-PvPLC3 | 591 | 67.32 | 5.73 | 57.05 | -0.508 | Cytoplasmic |
| PI-PvPLC4 | 599 | 68.74 | 5.75 | 52.51 | -0.56 | Cytoplasmic |
| PI-PvPLC5 | 595 | 68.28 | 9.35 | 50.94 | -0.482 | Mitochondrial |
| PI-PvPLC6 | 605 | 69.82 | 6.13 | 45.98 | -0.607 | Cytoplasmic |
| PI-PvPLC7 | 572 | 65.87 | 7.17 | 42.2 | -0.559 | Nuclear |
| PvNPC1 | 520 | 58.52 | 8.25 | 43.28 | -0.401 | Mitochondrial |
| PvNPC2 | 525 | 58.82 | 6.39 | 36.37 | -0.295 | Cytoplasmic |
| PvNPC3 | 515 | 57.48 | 6.77 | 39.97 | -0.277 | Cytoplasmic |
| PvNPC4 | 516 | 58.44 | 5.53 | 45.98 | -0.497 | Cytoplasmic |
| PvNPC5 | 487 | 54.50 | 6.98 | 36.96 | -0.256 | Cytoplasmic |
