## Supplementary Figures for "The non-specific phospholipase C of common bean *PvNPC4* modulates roots and nodule development"

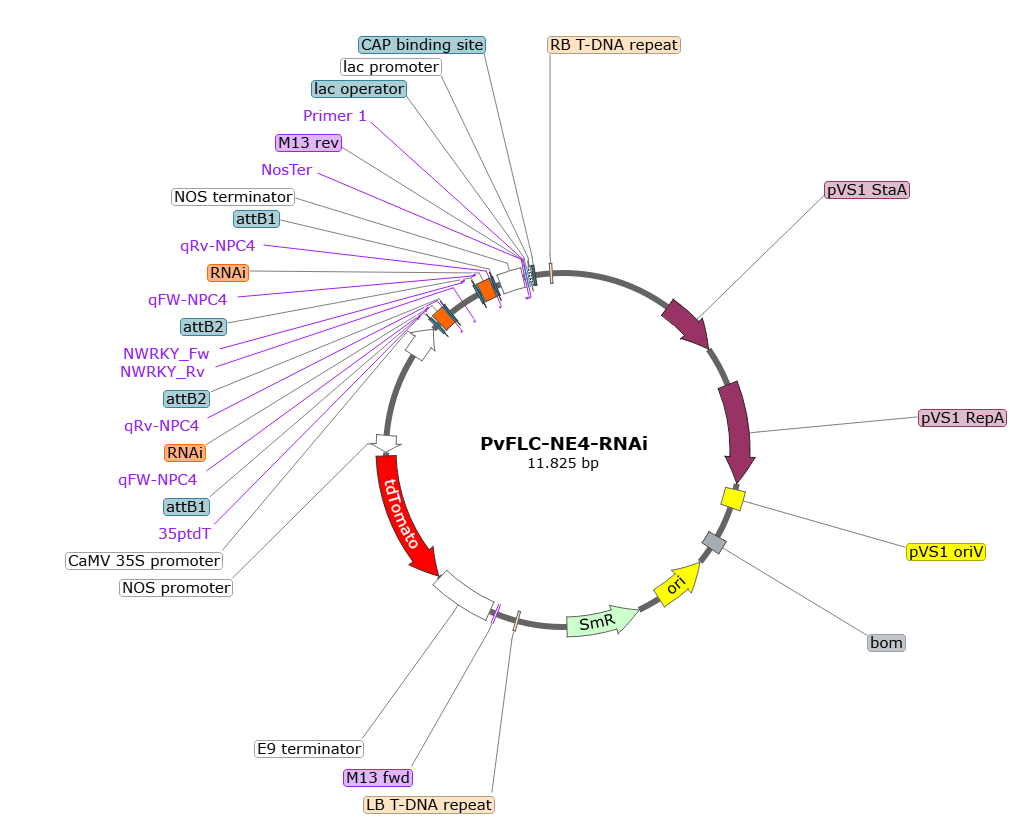

**S1 Fig.** ***In silico* map of the *PvNPC4*-RNAi vector.** This plasmid contains a double insertion of the RNAi fragment. The image was created with version 7.1.1 of SnapGene software.

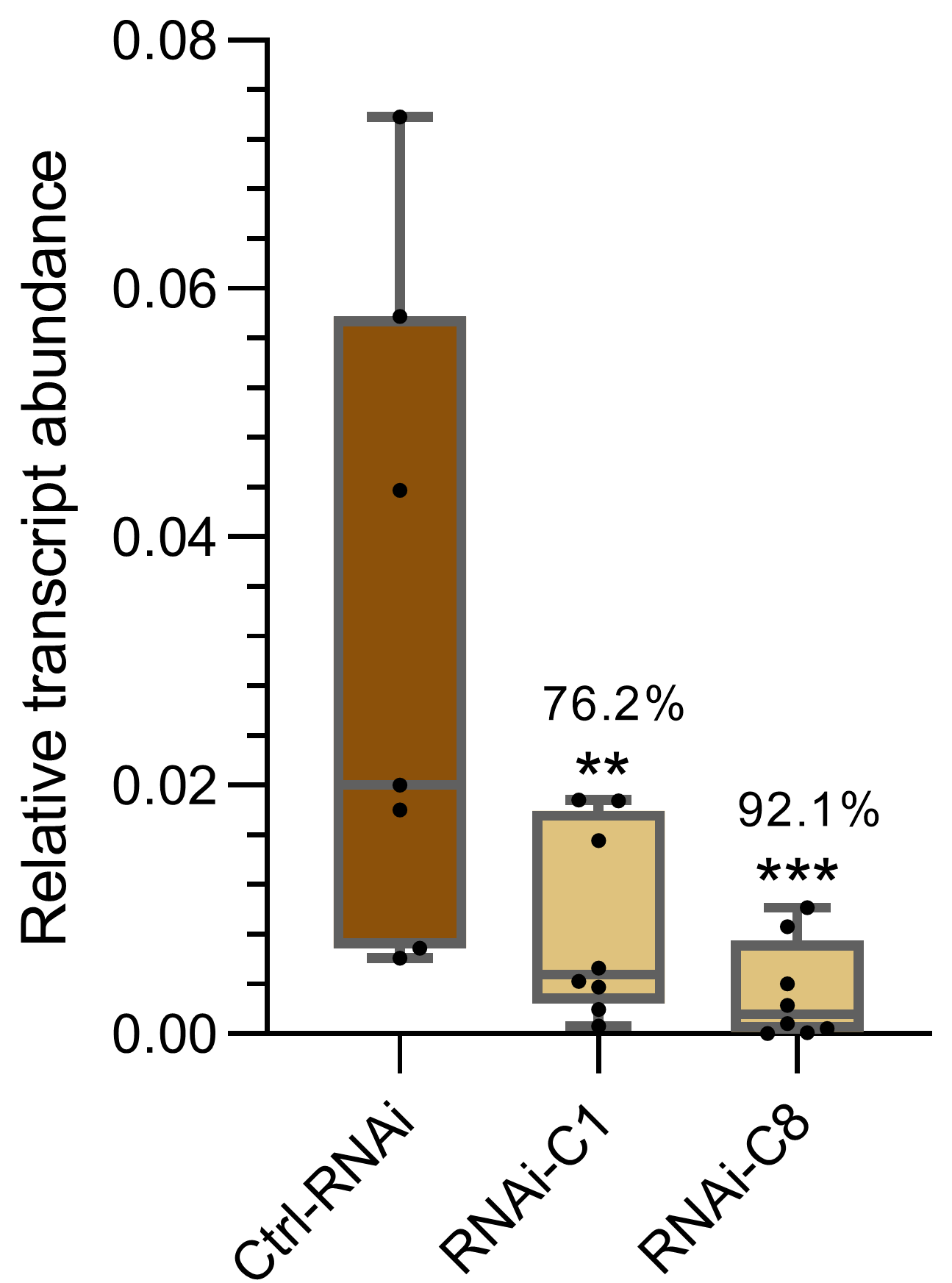

**S2 Fig. *PvNPC4* transcripts abundance in hairy roots carrying the control vector (pTdT-SAC) and the *PvNPC4*-RNAi silencing vector (RNAi-C1 and RNAi-C8 lineas)**. RNA-C1, *A. rhizognes* clone 1, which reduces *PvNPC4* transcript abundance in a 76,2%. RNA-C2, *A. rhizognes* clone 1, which reduces *PvNPC4* transcript abundance in a 92. 1%. The lower and upper edges of the boxes delimit the first to third quartiles, respectively, the central horizontal line represents the median, and the whiskers indicate the maximum and minimum values in the data set. Statistical significance was assessed with a Monte Carlo simulation test with 9999 resamples without replacement. (** *p* ≤ 0.01, *** *p* ≤ 0.001). Black dots in the box plots indicate independent samples from three biological replicates.

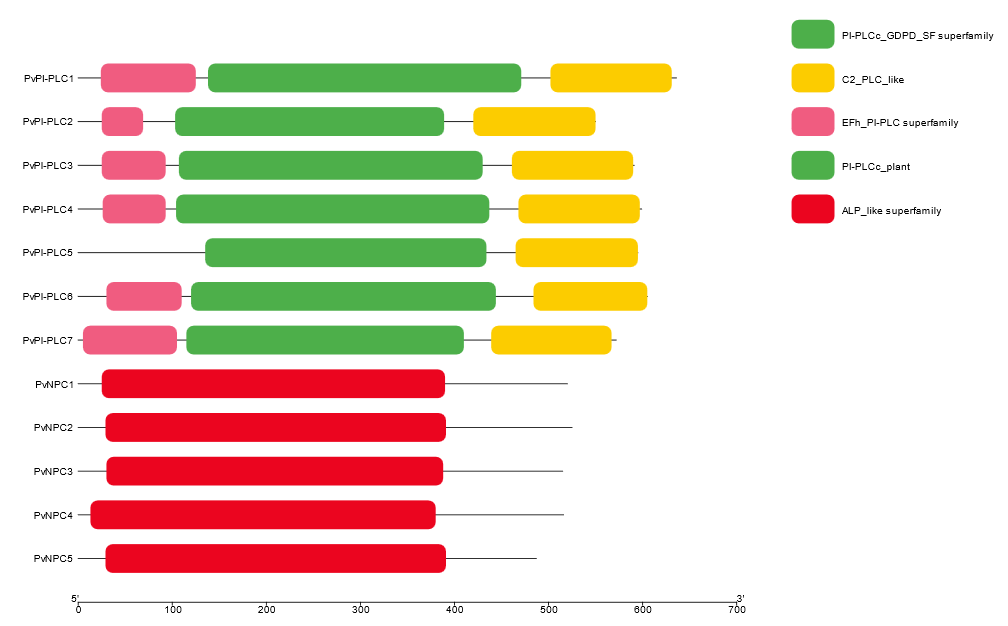

**S3 Fig.** **Distribution of conserved protein domains and motifs in PvPLC**. Pink box indicates the hand-like EF motif; green box indicates the PLC X and Y catalytic domains; yellow box indicates the C2 domain in PI-PLC. Red box indicates the phosphoesterase domain of the NPC subfamily.

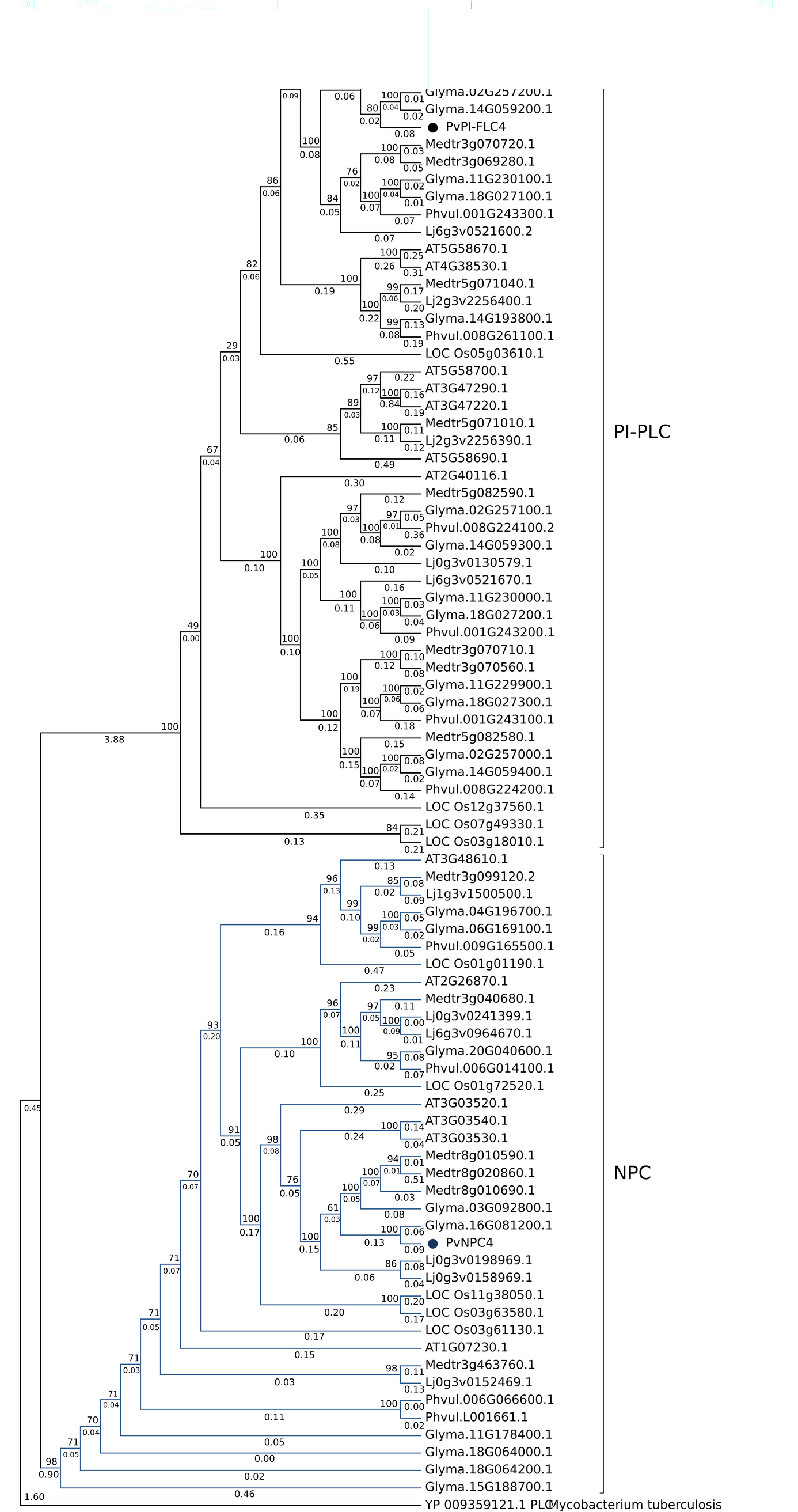

**S4 Fig. Phylogenetic tree inferred using the PLC amino acid sequences of leguminous and non-leguminous species.** *A. thaliana* (AT), *O. sativa* (LOC Os),
*L. japonicus*, *M. truncatula*, and *P. vulgaris* (Phvul). The black branches form the PI-PLC clade and the blue branches form the NPC clade. Black and blue circles indicate
PvPI-PLC4 and PvNPC, respectively. Decimal numbers on branches indicate the phylogenetic distance. The amino acid sequence of *M. tuberculosis* PLC (YP_009359121.1) was used as an external sequence. The phylogenetic tree was reconstructed using the IQ-TREE algorithm based on the maximum-likelihood method and the JTT+G model; 10,000 bootstraps were made.

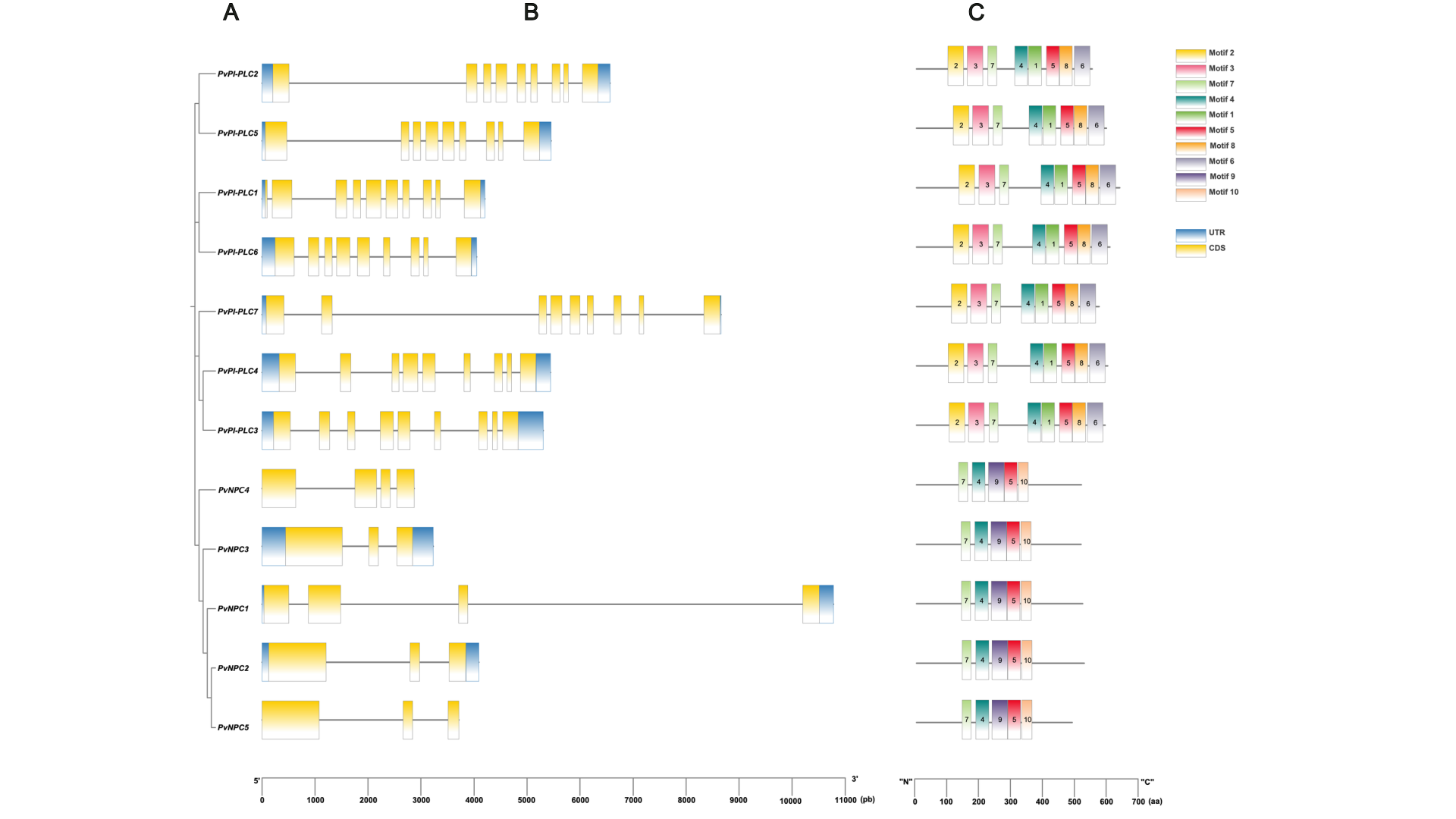

**S5 Fig. Analysis of the gene and protein sequence of PvPLC.** (A) Phylogenetic tree showing the relationship between the 12 PvPLC proteins. (B) Exon-intron structure of the 12 *PvPLC* genes. Yellow boxes represent exons and gray lines represent introns; green boxes represent UTR regions. (C) Motifs identified in PvPLC protein sequences using the MEME tool.

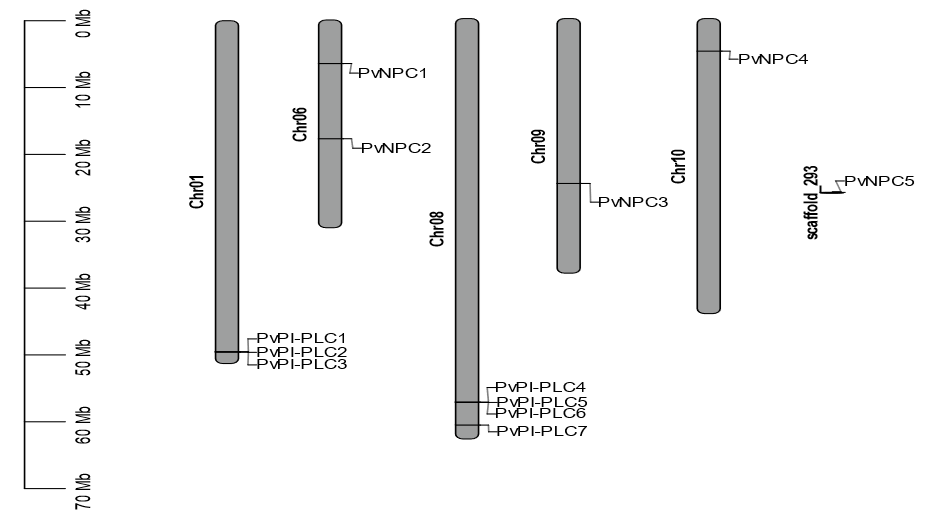

**S6 Fig.** Chromosomal location of *PvPLC* genes. The scale bar on the left indicates the chromosome sizes in megabase pairs (Mbp).

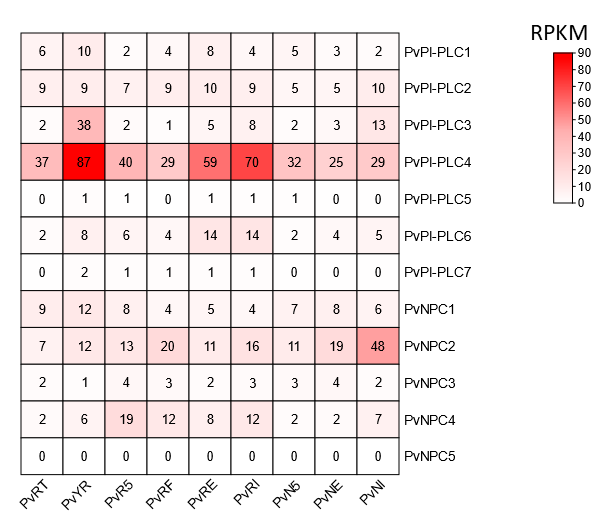

**S7 Fig. *In silico* PvGEA analysis showing the *PvPLCs* transcript abundance (RPKM) in different roots tissues and nodules**. PvRT: Root tips, 0.5-cm tissue, collected from fertilized plants at 2nd trifoliate stage of development. PvYR: Whole roots, including root tips, collected in the 2nd trifoliate stage of development. PvR5: Whole roots separated from 5-day old pre-fixing nodules. PvRF: Whole roots of fertilized plants collected at the same time as RE and RI. PvRE: Whole roots separated from fix+ nodules collected 21 days after inoculation. PvRI: Whole roots separated from fix-nodules collected 21 days after inoculation. PvN5: Pre-fixing (effective) nodules collected 5 days after inoculation. PvNE: Effectively fixing nodules collected 21 days after inoculation. PvNI: Ineffectively fixing nodules collected 21 days after inoculation.

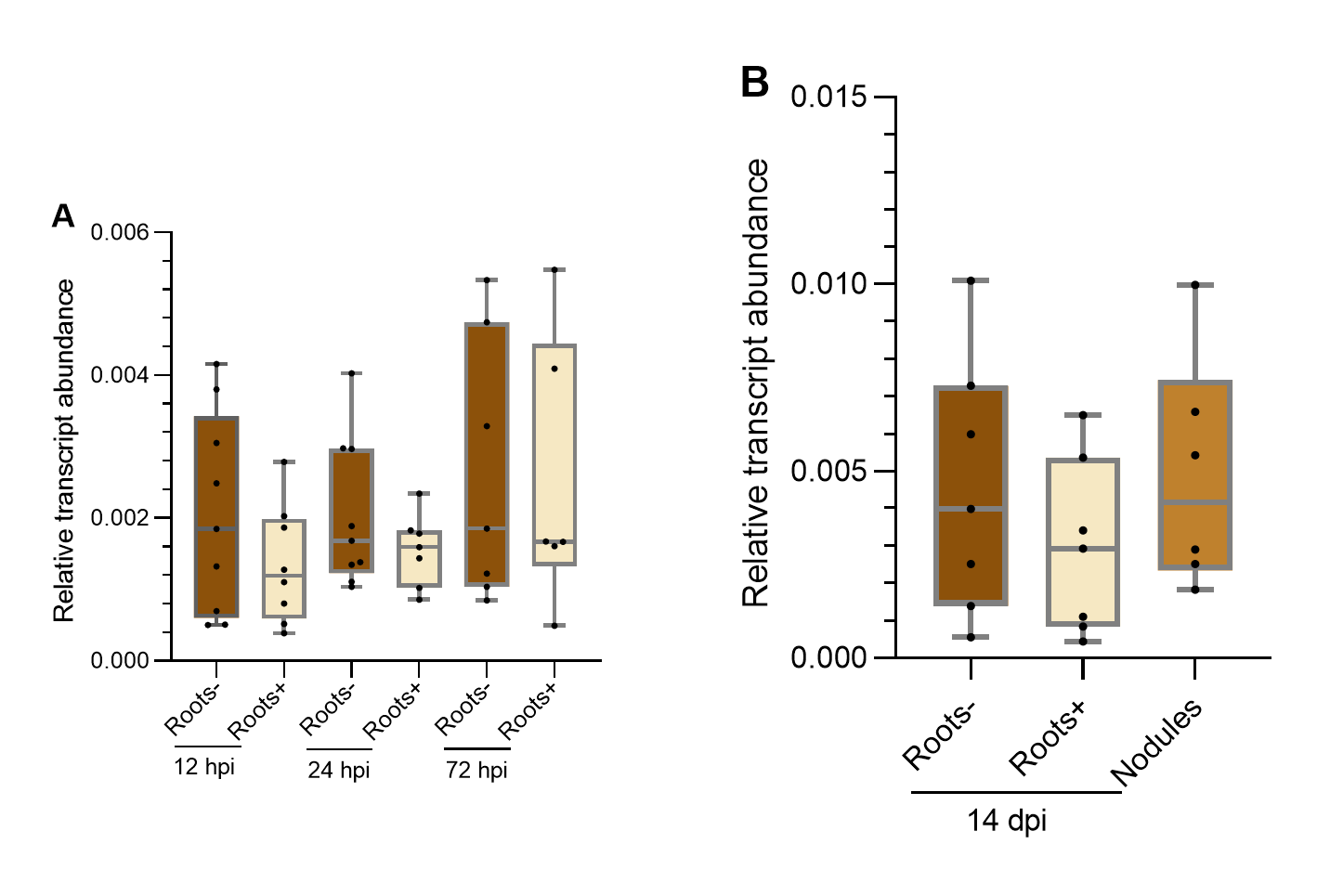

**S8 Fig.** ***R. tropici*** **does not affect *PvPI-PLC4* transcript abundance in wild-type roots and nodules of common bean**. (A) Early stages after *R. tropici* inoculation. (B) 14 dpi with *R. tropici*. Roots-, non-inoculated roots; Roots+, inoculated roots. The lower and upper edges of the boxes delimit the first to third quartiles, respectively, the central horizontal line represents the median, and the whiskers indicate the maximum and minimum values in the data set. Statistical significance was assessed with a Monte Carlo simulation test with 9999 resamples without replacement. Black dots in the box plots indicate independent samples from three biological replicates.

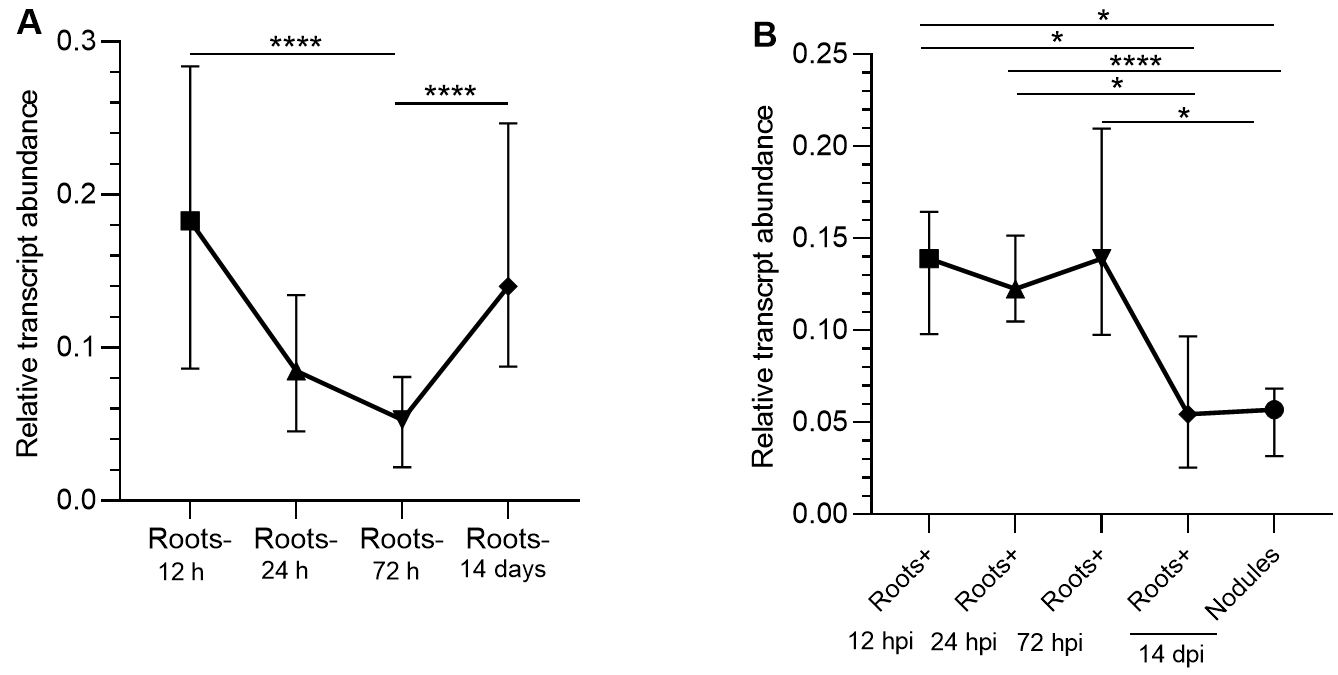

**S9 Fig. *In silico* analysis *PvNPC4* transcript abundance fluctuates in common bean roots according to the stage of development**. (A) Non-inoculated roots (Root-). (B) Roots inoculated with *R. tropici* (Roots-). For the statistical analysis a Monte Carlo simulation test was performed with 9999 resamples without replacement. The geometric shapes of the trend line indicate the median and the whiskers indicate the interquartile range. Three biological replicates were performed (* *p* ≤ 0.05, **** *p* ≤ 0.0001).

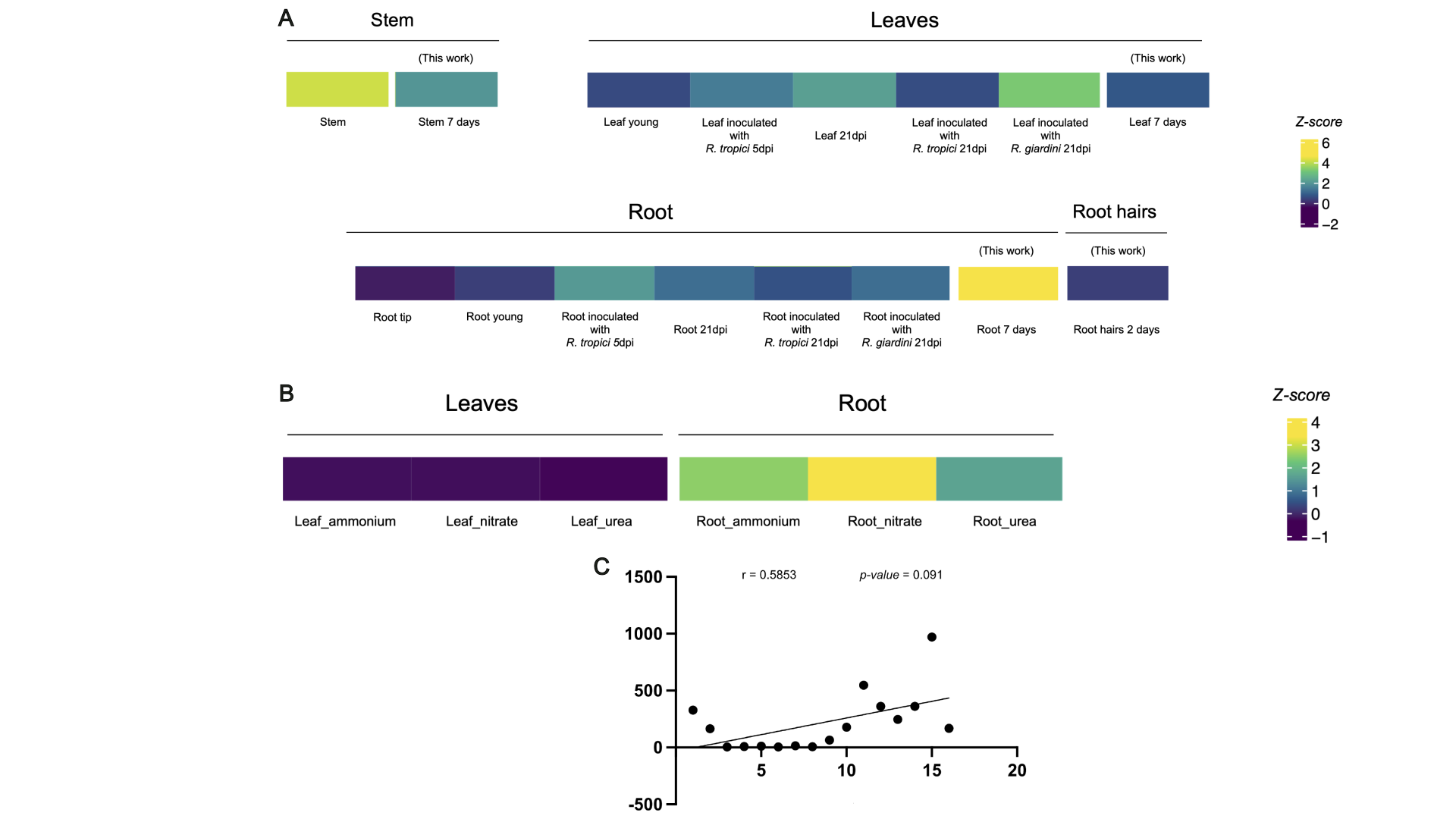

**S10 Fig.** ***PvNPC4* transcriptional landscape in wild-type common bean tissues**.
(A) *PvNPC4* transcript abundance patterns in different tissues and developmental stages of *P. vulgaris* inoculated with *R. tropici* or *Rhizobium giardini* and in non-inoculated plants at 7 days after transplanting into pots (this work). (B) *AtNPC4* transcriptional landscape in leaf and root under different conditions. (C) Spearman correlation between *PvNPC4* transcript abundance in common bean tissues and its expression obtained by RNA-seq. Z-score values are represented in fold change (FC).
