## Supplementary material for "The non-specific phospholipase C of common bean *PvNPC4* modulates roots and nodule development": Heatmap

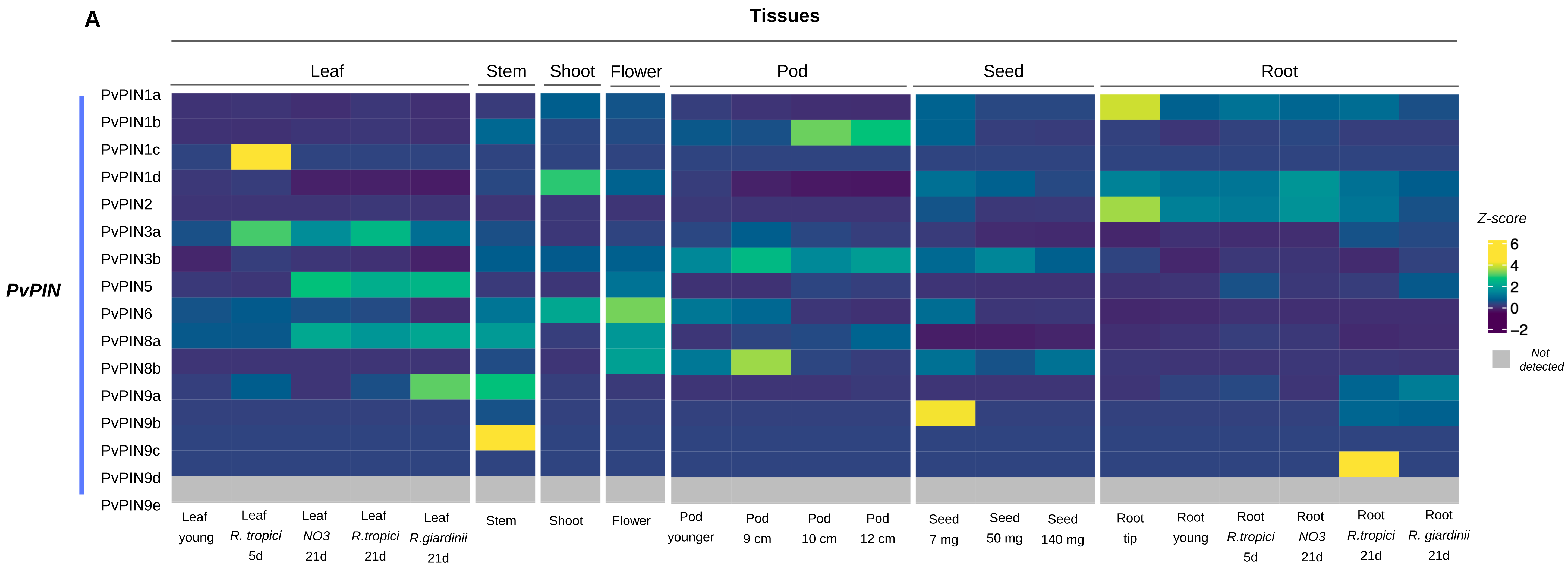

A
Tissues
Leaf
Stem
Shoot
Flower
Pod
Seed
Root
PvPIN
PvPIN1a
PvPIN1b
PvPIN1c
PvPIN1d
PvPIN2
PvPIN3a
PvPIN3b
PvPIN5
PvPIN6
PvPIN8a
PvPIN8b
PvPIN9a
PvPIN9b
PvPIN9c
PvPIN9d
PvPIN9e
Leaf young
Leaf R. tropici 5d
Leaf NO3 21d
Leaf R.tropici 21d
Leaf R.giardinii 21d
Stem
Shoot
Flower
Pod younger
Pod 9 cm
Pod 10 cm
Pod 12 cm
Seed 7 mg
Seed 50 mg
Seed 140 mg
Root tip
Root young
Root R.tropici 5d
Root NO3 21d
Root R.tropici 21d
Root R. giardinii 21d
Z-score
6
4
2
0
−2
Not detected

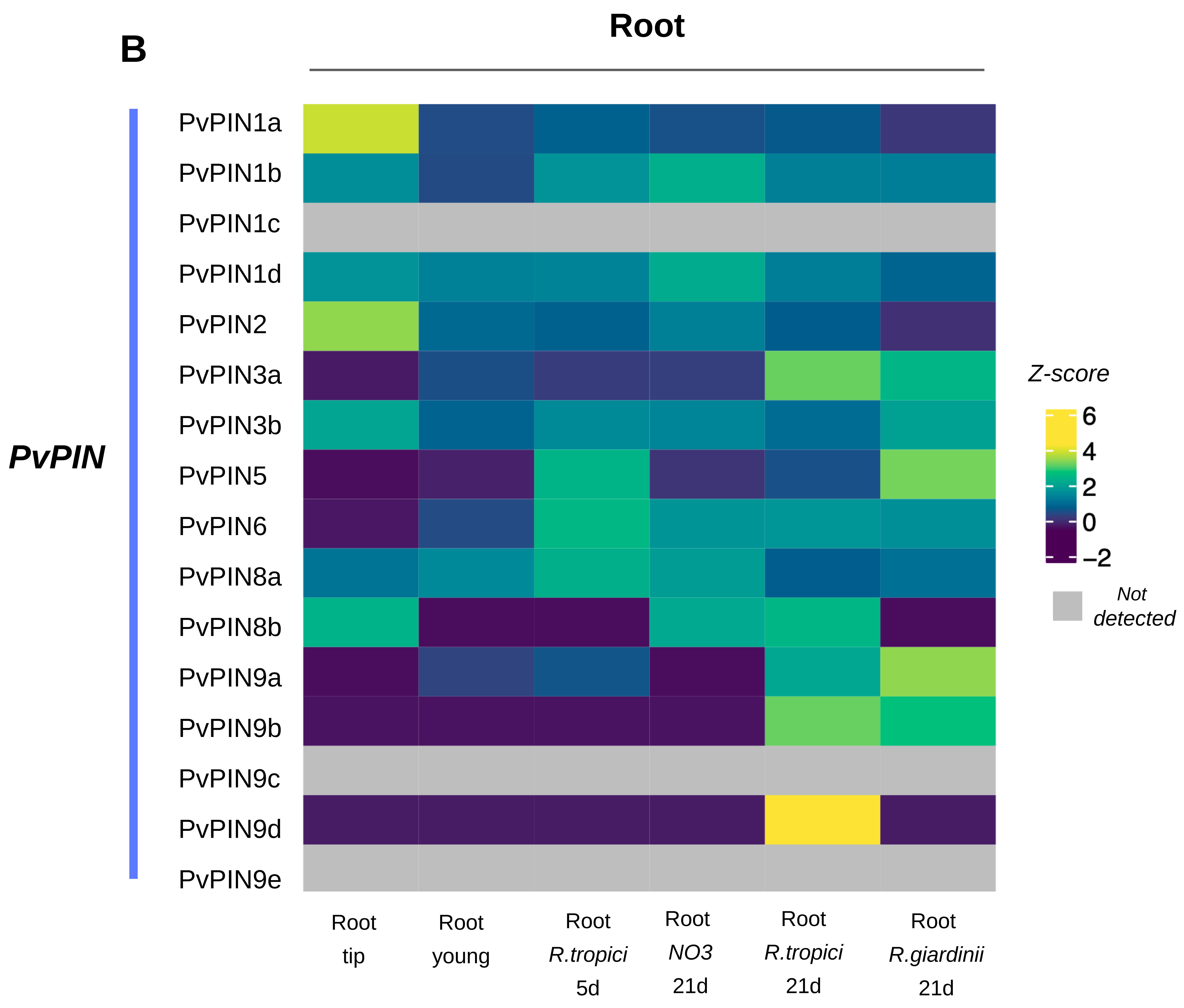

B
Root
PvPIN
PvPIN1a
PvPIN1b
PvPIN1c
PvPIN1d
PvPIN2
PvPIN3a
PvPIN3b
PvPIN5
PvPIN6
PvPIN8a
PvPIN8b
PvPIN9a
PvPIN9b
PvPIN9c
PvPIN9d
PvPIN9e
Root tip
Root young
Root R.tropici 5d
Root NO3 21d
Root R.tropici 21d
Root R.giardinii 21d
Z-score
6
4
2
0
−2
Not detected

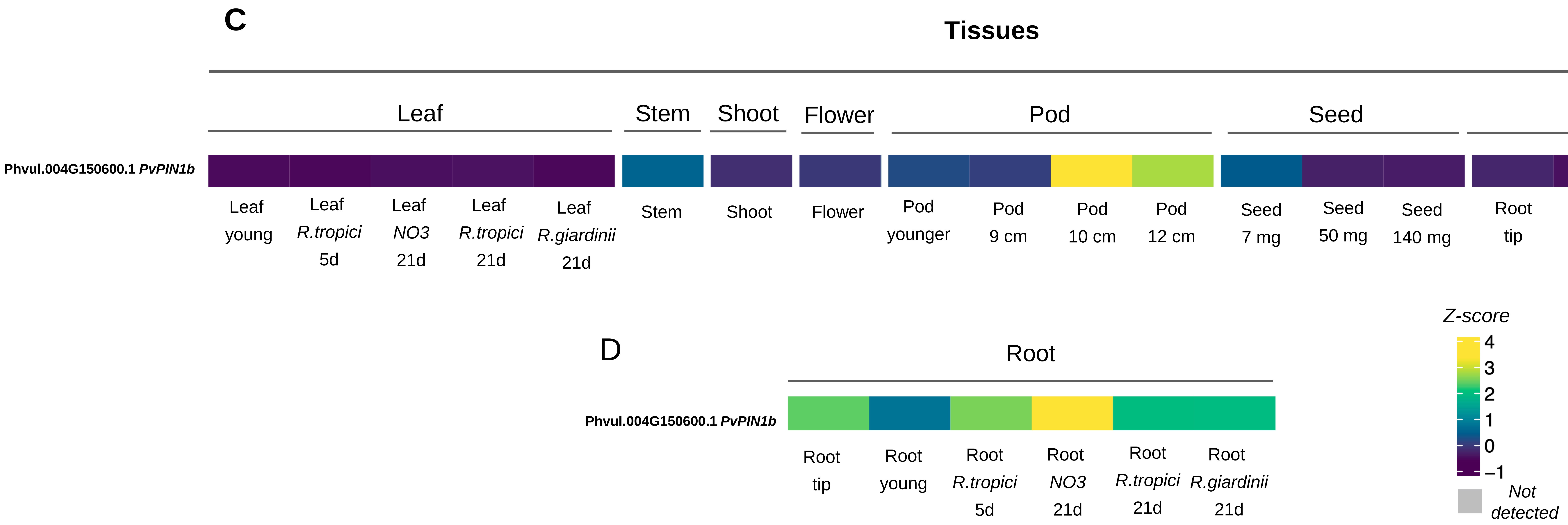


**S1 Fig. Transcriptional landscape of *PvPin* genes in wild-type common bean tissues**. (A) Transcriptional landscape of *PvPin* genes in different tissues (PvGEA data, RPKM). (B) Transcriptional landscape of *PvPin* genes in different root tissue treatments (Open Big Data metatranscriptome). (C) Transcriptional landscape of *PvPin1b* in different tissues (PvGEA data, RPKM). (D)Transcriptional landscape of *PvPin1b* in different root tissue treatments (Open Big Data metatranscriptome).

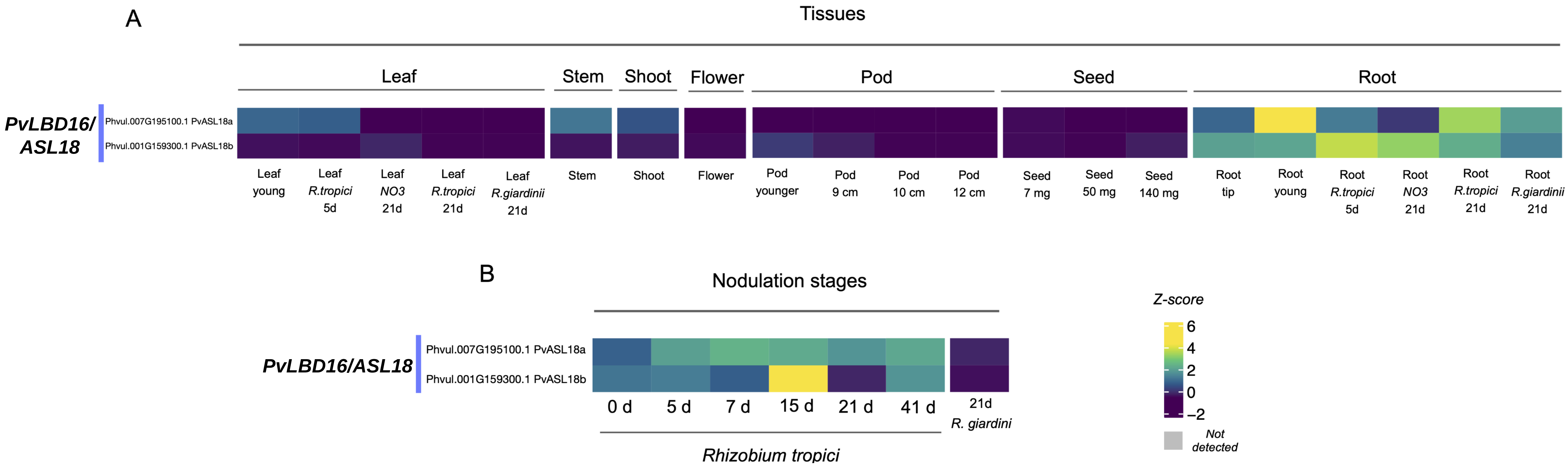


**S2 Fig. Transcriptional landscape of *PvASL18a* and *ASL18b* genes in wild-type common bean tissues**. (A) Transcriptional landscape in different tissues (PvGEA data, RPKM). (B) Transcriptional landscape in different root tissue treatments and nodules (Open Big Data metatranscriptome ).

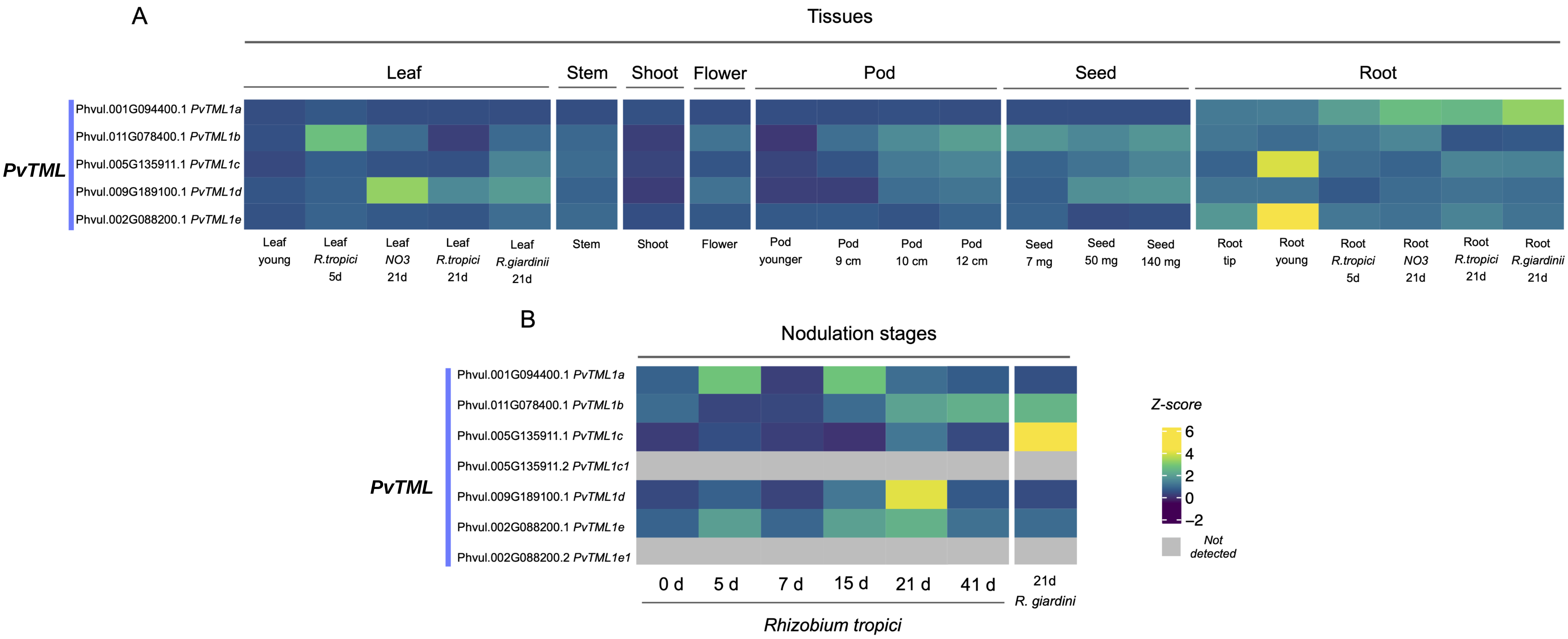


**S3 Fig. Transcriptional landscape of *PvTML* genes in wild-type common bean tissues**. (A) Transcriptional landscape in different tissues (PvGEA data, RPKM). (B) Transcriptional landscape in different root tissue treatments and nodules (Open Big Data metatranscriptome).
